# Mean-Variance QTL Mapping Identifies Novel QTL for Circadian Activity and Exploratory Behavior in Mice

**DOI:** 10.1101/276972

**Authors:** Robert W. Corty, Vivek Kumar, Lisa M. Tarantino, Joseph S. Takahashi, William Valdar

**Author notes:** Corresponding author: William Valdar, 120 Mason Farm Road, Genetic Medicine Building, Suite 5113, Campus Box 7264, Chapel Hill, NC, 27599, USA.

## Abstract

We illustrate, through two case studies, that “mean-variance QTL mapping” can discover QTL that traditional interval mapping cannot. Mean-variance QTL mapping is based on the double generalized linear model, which elaborates on the standard linear model by incorporating not only a linear model for the data itself, but also a linear model for the residual variance. Its potential for use in QTL mapping has been described previously, but it remains underutilized, with certain key advantages undemonstrated until now. In the first case study, we use mean-variance QTL mapping to reanalyze a reduced complexity intercross of C57BL/6J and C57BL/6N mice examining circadian behavior and find a mean-controlling QTL for circadian wheel running activity that was not detected by traditional interval mapping. Mean-variance QTL mapping was more powerful than traditional interval mapping at the QTL because it accounted for the fact that mice homozygous for the C57BL/6N allele had less residual variance than the other mice. In the second case study, we reanalyze an intercross between C57BL/6J and C58/J mice examining anxiety-like behaviors, and identify a variance-controlling QTL for rearing behavior. This QTL was not identified in the original analysis because traditional interval mapping does not target variance QTL.

## INTRODUCTION

Interval mapping based on the assumption of homogeneous residual variance and its associated statistical methods (Lander and Botstein 1989; Martínez and Curnow 1992; Haley and Knott 1992; Churchill and Doerge 1994) have successfully identified many QTL for important complex traits over the last 25 years. It is well-appreciated, however, that this focus on mean effects fails to capture all the complexities of the relationship between genotype and phenotype, and recent methodological developments have sought to expand the repertoire of detectable patterns of association (Geiler-Samerotte *et al.* 2013; Nelson *et al.* 2013; Shen *et al.* 2012; Forsberg *et al.* 2015; Paré *et al.* 2010).

Two closely related developments target genotypically-induced differences in the residual variance, while simultaneously controlling for possible effects on the mean: the application of the double generalized linear model (DGLM; Smyth 1989) to QTL mapping (Rönnegård and Valdar 2011) and the omnibus test of Cao *et al.* (2014) for genetic heterogeneity. Both of these approaches can detect standard QTL influencing phenotype mean (mQTL), QTL influencing phenotype variance (vQTL) and QTL influencing some combination of phenotype mean and variance (mvQTL) (see Corty and Valdar 2018+ [BVH]). Despite the demonstrated potential of these methods to detect QTL that current standard methods overlook, they remain underutilized.

Barriers to widespread adoption include a lack of proven potential in real data applications and the absence of software that is interoperable with existing infrastructure. Apart from those barriers, one reasonable concern is that a novel approach might fail to identify known QTL, adding needless complexity to the interpretation of already-reported QTL. This concern should be largely allayed by the nature of the DGLM as an extension of the linear model, simplifying to the latter when variance heterogeneity is absent. In fact, rather than add complexity, the DGLM automatically classifies QTL into mQTL, vQTL, or mvQTL, clarifying the genotype-phenotype relationship.

Here we demonstrate, with two real data examples available from the Mouse Phenome Database (Bogue *et al.* 2015), that QTL mapping using the DGLM, which we term “mean-variance QTL mapping” largely replicates the results of standard QTL mapping and detects additional QTL that the traditional analysis does not. In two companion articles, we demonstrate typical usage of `R` package `vqtl`, which implements mean-variance QTL mapping (Corty and Valdar 2018+ [r/vqtl]) and describe the unique ability of mean-variance QTL mapping and its associated permutation procedure to reliably detect QTL in the face of variance heterogeneity that arises from non-genetic factors (Corty and Valdar 2018+ [BVH]).

## STATISTICAL METHODS

### Traditional QTL mapping based on the standard linear model (SLM)

The traditional approach to mapping a quantitative trait in an experimental cross with no population structure (*e.g.* an F2 intercross or backcross) involves fitting, at each putative locus in turn, a linear model of the following form. Letting *y*_*i*_ denote the phenotype value of individual *i*, this phenotype is modeled as

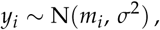

where *σ*^2^ is the residual variance, and the expected phenotype mean, *m*_*i*_, is predicted by effects of QTL genotype and, optionally, effects of covariates. In the reanalyses performed here, *m*_*i*_ is modeled to include a covariate of sex and additive and dominance effects of QTL genotype, that is,

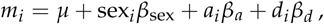

where *µ* is the intercept, *β*_sex_ is the sex effect, with sex_*i*_ indicating (0 or 1) the sex of individual *i*, and *β*_*a*_ and *β*_*d*_ are the additive and dominance effects of a QTL whose genotype is represented by *a*_*i*_ and *d*_*i*_ defined as follows: when QTL genotype is known, *a* _*i*_ is the count (0,1,2) of one parental allele, and *d*_*i*_ indicates heterozygosity (0 or 1); when QTL genotype is inferred based on flanking marker data, as is done here, *a*_*i*_ and *d*_*i*_ are replaced by their corresponding probabilistic expectations (Haley and Knott 1992; Martínez and Curnow 1992). The evidence for association at a given putative QTL is based on a comparison of the fit of the model above with that of a null model that is identical except for the QTL effects being omitted. These models and their comparison we henceforth refer to as the standard linear model (SLM) approach.

### Mean-variance QTL mapping based on the double generalized linear model (DGLM)

The statistical model underlying mean-variance QTL mapping, the double generalized linear model (DGLM; Smyth 1989 and Rönnegård and Valdar 2011), elaborates the SLM approach by modeling a potentially unique value of *σ*^2^ for each individual, as

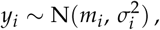

where *m*_*i*_ has the same meaning as in the SLM, but now 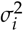 is linked to its own linear predictor *v*_*i*_ as

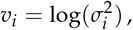

where the logarithm ensures that 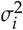 is always positive. The linear predictors for *m*_*i*_ and *v*_*i*_ are modeled as

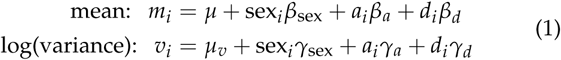

where *µ, a*_*i*_, *d*_*i*_, sex_*i*_, and the *β*’s are as before, *µ*_*v*_ is an intercept representing the (log of the) “baseline” residual variance, and *γ*_*a*_, *γ*_*d*_, and *γ*_sex_ are the effects of the QTL and covariates on *v*_*i*_.

The evidence for a QTL association is now defined through three distinct model comparisons, corresponding to testing for an mQTL, a vQTL, or an mvQTL. In each case, the fit of the “full” model in Equation 1 is compared with that of a different fitted null: for the mQTL test, the null model omits the QTL effects on the mean (*i.e., β*_*a*_ = *β*_*d*_ = 0); for the vQTL test, the null model omits the QTL effects on the variance (*i.e., γ*_*a*_ = *γ*_*d*_ = 0); and for the mvQTL test, the null model omits QTL effects on both mean and variance (*i.e., β*_*a*_ = *β*_*d*_ = *γ*_*a*_ = *γ*_*d*_ = 0).

### Genomewide significance and FWER-adjusted p-values

The model comparisons described for both the SLM test and the three DGLM-based tests each produce likelihood ratio (LR) statistics. These LR statistics are converted to *p*-values that are adjusted for the family-wise error rate (FWER) across loci, *i.e., p*-values on the scale of genomewide significance. This adjustment is performed separately for each test by calculating an empirical distribution for the LR statistic under permutation, much in the spirit of Churchill and Doerge (1994) but with some modifications, namely that different tests have differently structured permutations. Briefly, let *G*_*i*_ be the full set of genetic information for individual *i*, that is, the genotypes or genotype probabilities across all loci. For the SLM and mvQTL tests, we define a permutation as randomly shuffling the *G*_*i*_ ‘s across individuals; for the mQTL test, the permutations apply this shuffle only to the genotype information in the full model’s mean component; for the vQTL test, the permutations apply the shuffle only to the genotype information in the full model’s variance component. For a given test, for each permutation we calculate LR statistics across the genome and record the maximum; the maxima of over all permutations is fitted to a generalized extreme value distribution, and the upper tail probabilities of this fitted distribution are used to calculated the FWER-adjusted *p*-values for the LR statistics in the unpermuted data [see Dudbridge and Koeleman 2004, and, *e.g.*, Valdar *et al.* 2006; more details in Corty and Valdar 2018+ [BVH]]. An FWER-adjusted *p*-value can be interpreted straightforwardly: it is the probability of observing an association statistic this large or larger in a genome scan of a phenotype with no true associations.

### Data Availability and Software

Supplementary file S1 contains the dataset from Kumar *et al.* (2013), also available from the Mouse Phenome Database (Bogue *et al.* 2015) at https://phenome.jax.org/projects/Kumar1. Supplementary file S2 contains the script to run the genome scans presented here. Supplementary file S3 contains the script to plot the results of that analysis. Supplementary file S4 contains the dataset from Bailey *et al.* (2008), also available from the Mouse Phenome Database at https://phenome.jax.org/projects/Bailey1. Supplementary file S5 contains the script to run the genome scans presented here. Supplementary file S 6 contains the script to plot the results of that analysis. Supplementary file S 7 contains the script to trim out redundant information from the results to make them easier to share online. Supplementary file S8 contains the script that runs the power simulation comparing the DGLM to the SLM at the QTL identified in the Kumar reanalysis.

The replications of the original analyses were conducted with `R` package `qtl` and reanalyses with mean-variance QTL mapping were conducted with `R` package `vqtl`, both freely available on CRAN. A companion article demonstrates typical usage of package `vqtl` (Corty and Valdar 2018+ [r/vqtl]).

## REANALYSIS OF KUMAR ET AL. REVEALS A NEW MQTL FOR CIRCADIAN WHEEL RUNNING ACTIVITY

### Summary of Original Study

Kumar *et al.* (2013) intercrossed C57BL/6J and C57BL/6N, two closely-related strains of C57BL6 that diverged in 1951, approximately 330 generations ago. Due to recent coancestry of the parental strains, this cross is termed a “reduced complexity cross”, and their limited genetic differences ensure that any identified QTL region can be narrowed to a small set of variants bioinformatically. The intercross resulted in 244 F2 offspring, 113 female and 131 male, which were tested for acute locomotor response to cocaine (20mg/kg) in the open field. One to three weeks following psychostimulant response testing, the mice were tested for circadian wheel running activity.

Analysis of wheel running data was carried out using Clock-Lab software v6.0.36. For calculation of activity 20 day epoch in DD was used in order to have standard display between actograms. Analysis of other circadian measures such as period (tau) or amplitude were carried out using methods previously described (Shimomura *et al.* 2001). All animal protocols were approved by the Institutional Animal Care and Use Committee (IACUC) of the University of Texas Southwestern Medical Center

Traditional QTL mapping with the SLM, reported in Kumar *et al.* (2013), detected a single large-effect QTL for cocaine-response traits on chromosome 11, but no QTL for circadian activity. A later study by another group nonetheless observed that the circadian activity of the two strains showed significant differences (Banks *et al.* 2015).

### Reanalysis with traditional QTL mapping and mean-variance QTL mapping

For the cocaine response traits, traditional QTL mapping and mean-variance QTL mapping gave results that were nearly identical to the originally-published analysis in Kumar *et al.* (2013) (Figure S1).

For the circadian wheel running activity trait, however, traditional QTL mapping identified no QTL (Figure 1 in green) but mean-variance QTL mapping identified one QTL on chromosome 6 (Figure 1 in blue, black, and red). In this case, all three tests were statistically significant, but the most significant was the mQTL test (blue), so we discuss it as an mQTL. The most significant genetic marker was rs30314218 on chromosome 6, at 18.83 cM, 40.0 Mb, with a FWER-controlling *p*-value of 0.0063. The mQTL explains 8.4% of total phenotype variance by the traditional definition of percent variance explained (*e.g.*, Broman and Sen 2009).

**Figure 1.**
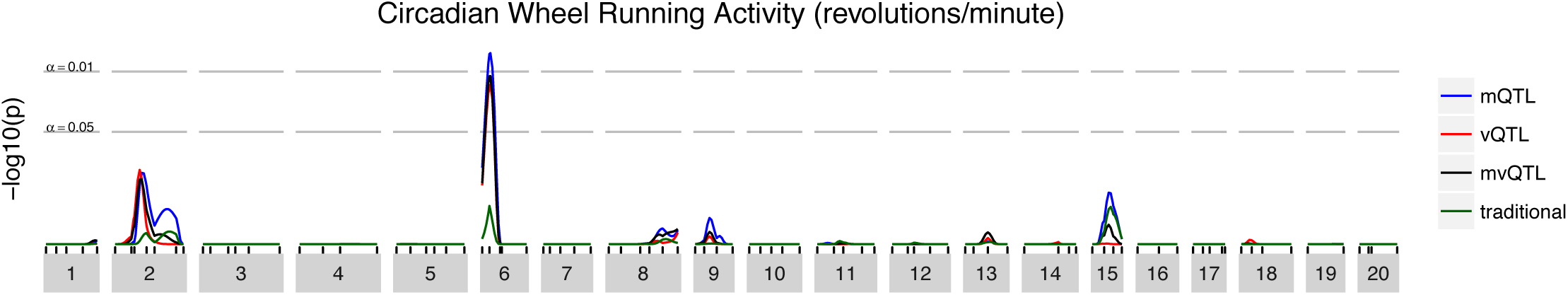
Genome scan for Kumar et al. circadian wheel running activity. The horizontal axis shows chromosomal location and the vertical axis shows FWER-controlling *p*-values for the association between each genomic locus and circadian wheel running activity.

### Understanding the Novel QTL

Though they test for the same pattern, the mQTL test of mean-variance QTL mapping identified a QTL where the traditional QTL test did not. This discordance may arise when there is variance heterogeneity in the mapping population. In this case, mice homozygous for the C57BL/6N allele at the mQTL have both higher average wheel running activity and lower residual variance in wheel running activity than mice with other genotypes (Figure 2a).

**Figure 2.**
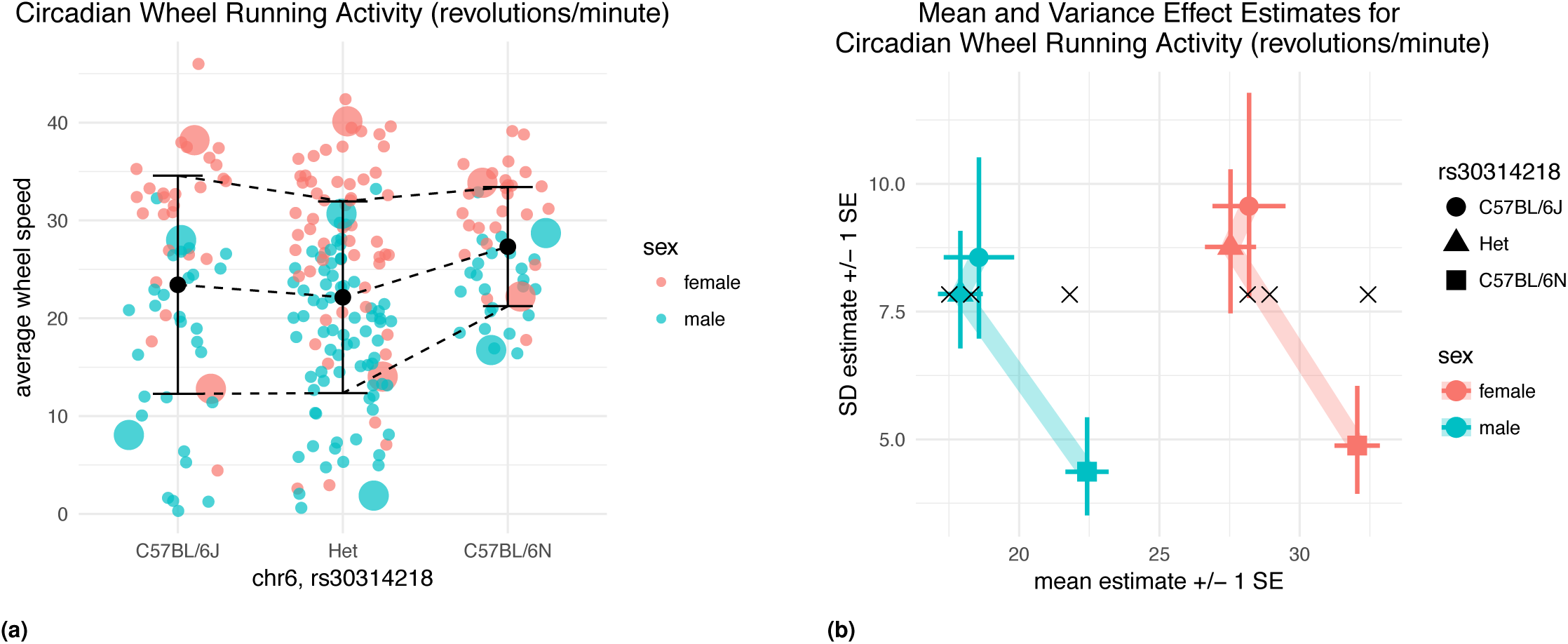
(a) Average wheel speed (revolutions/minute) of all mice. It is visually apparent that female mice had higher circadian wheel running activity than male mice and that mice that homozygous for C57BL/6N had higher circadian wheel running activity and less intra-genotype variation. Large dots indicate the mice whose activity is shown in actogram form (Figure 3). (b) Predicted mean and variance of mice according to sex and allele at the QTL. What was visually apparent in (a) is captured by the DGLM. The estimated parameters relating to mice that are homozygous for the C57BL/6N allele imply a higher expected value and a lower residual variance than the other two genotype groups. Black x’s indicate the estimates from the SLM, very similar to the DGLM estimates in the horizontal (mean) axis, but homogeneous in the vertical (variance) axis.

The identification of this QTL by mean-variance QTL mapping but not traditional QTL mapping can be understood by contrasting how the DGLM and SLM fit the data at this locus.

For the SLM, a single value of the residual standard deviation *σ* is estimated for all mice. Approximately 25% of the mice are homozygous for the C57BL/6N allele, so *σ* is estimated mostly based on heterozygous mice and homozygous C57BL/6J mice. The SLM estimates 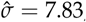, a slight underestimate for some genotype-sex combinations, and a drastic overestimate for the homozygous C57BL/6N of both sexes (Figure 2b). With *σ* overestimated for the C57BL/6N homozygotes, the addition of a locus effect to the null model results in only a limited increase in the likelihood. For the DGLM, six different values of *σ* are estimated, one for each genotype-sex combination (Figure 2b). In light of the better-estimated (lower) 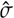 for the C57BL/6N homozygotes, the estimated mean effects are very unlikely due to chance alone.

A simulation based on the estimated coefficients shows that at a false positive rate of 5 × 10^*-*4^, relevant for genome-wide significance testing, the SLM has 63% power to reject the null at this locus and the DGLM has 91% power (Supplementary file S8).

### Variant Prioritization

Reduced complexity crosses allow variant prioritization to proceed quickly because of the number of segregating variants are few. Using 1000 nonparametric bootstrap resamples, the QTL interval was estimated as 13.5-23.5 cM (90% CI), which translates to physical positions of 32.5 - 48.5 Mb using Mouse Map Converter’s sex averaged Cox map (Cox *et al.* 2009). Since this interval contains no genes or previously identified QTL shown to regulate circadian rhythms, we prioritized candidates by identifying variants between C57BL/6J and C57BL/6NJ based on Sanger mouse genome database (Keane *et al.* 2011; Simon *et al.* 2013), which yielded 463 SNPs, 124 indels, and 3 structural variants (Table 1).

**Table 1.** Genetic Variants in QTL interval for circadian wheel running activity

| location | indel | SNP | SV | Total |
| --- | --- | --- | --- | --- |
| exon, missense | – | 2 | – | 2 |
| intron, splice region | 1 | – | – | 1 |
| intron, nonsynonymous | 57 | 246 | – | 303 |
| intron, synonymous | – | 1 | – | 1 |
| 3' UTR | – | 3 | – | 3 |
| upstream | 6 | 29 | – | 35 |
| downstream | 7 | 20 | – | 27 |
| intergenic | 53 | 161 | – | 214 |
| unclassified | – | 1 | 3 | 4 |

**Table 2.**
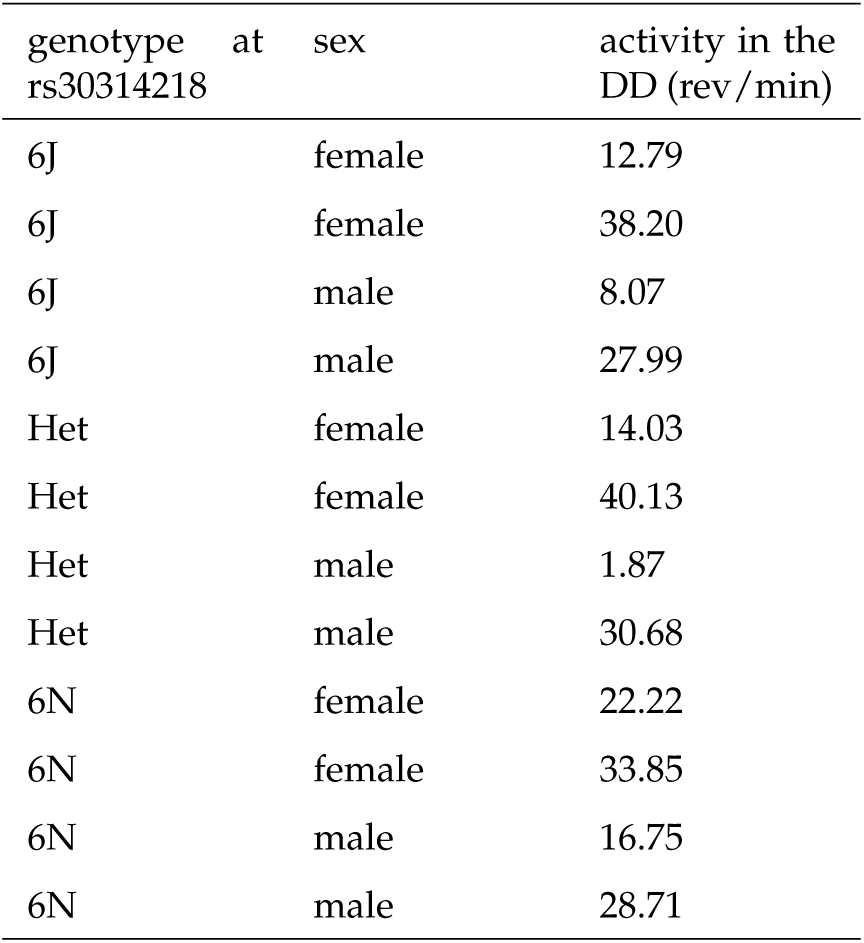
The characteristics of the mice plotted in Figure 3

Of these variants, none of the indels or structural variants were nonsynonymous. Two SNPs were predicted to lead to missense changes (T to A at position 6: 39400456 in *Mkrn1*, and A to A/C at 6:48486716 in *Sspo*). The variant in Sspo was a very low confidence call and therefore likely a false positive.

The *Mkrn1* (makorin, ring finger protein, 1) variant is a mutation in C57BL/6J and changes a highly conserved (Figure S2) tyrosine to asparagine and was determined to be the best candidate variant in the QTL interval. Mkrn1 is a ubiquitin E3 ligase with zinc fingers with poorly defined function (Kim *et al.* 2005). It is expressed at low levels widely in the brain according to Allen Brain Atlas and EBI Expression Atlas (Kapushesky *et al.* 2009; McWilliam *et al.* 2013; Allen Institute for Brain Science 2015; McWilliam *et al.* 2013). Functional analysis will be necessary to experimentally confirm that this variant in Mkrn1 is indeed the causative mutation that led, in a dominant fashion, to the decreased expected value and increased variance of circadian wheel running activity observed in mice with at least one copy of the C57BL/6J haplotype in the QTL region in this study.

## REANALYSIS OF BAILEY ET AL. IDENTIFIES A NEW VQTL FOR REARING BEHAVIOR

### Summary of Original Study

Bailey *et al.* (2008) intercrossed C57BL/6J and C58/J mice, two strains known to be phenotypically similar for anxiety-related behaviors, as a control cross for an ethylnitrosourea mutagenesis mapping study. The intercross resulted in 362 F2 offspring, 196 females and 166 males. Six open-field behaviors were measured at approximately 60 days of age in a 43cm by 43cm by 33cm white arena for ten minutes. All phenotypes were transformed with the rank-based inverse normal transform to limit the influence of outliers. The authors reported 7 QTL spread over five of the six measured traits, but none for rearing behavior.

### Reanalysis with SLM and DGLM

SLM-based QTL analysis replicated the originally-reported LOD curves. Significance thresholds to control FWER at 0.05 were estimated by 10,000 permutations, using the method described in the original publication, but found to be meaningfully higher than the originally-reported thresholds. Of the 7 originally-reported QTL, 3 exceeded the newly-estimated thresholds (Figure S4).

The DGLM-based reanalysis was initially conducted with the rank-based inverse normal transformed phenotypes, to maximize the comparability with the original study. This reanalysis largely replicated the results of the SLM-based analysis identified a statistically-significant vQTL for rearing behavior on chromosome 2 (Figure 4 and Figure S4). The top marker under the peak was at 38.6cM and 65.5Mb.

**Figure 3.**
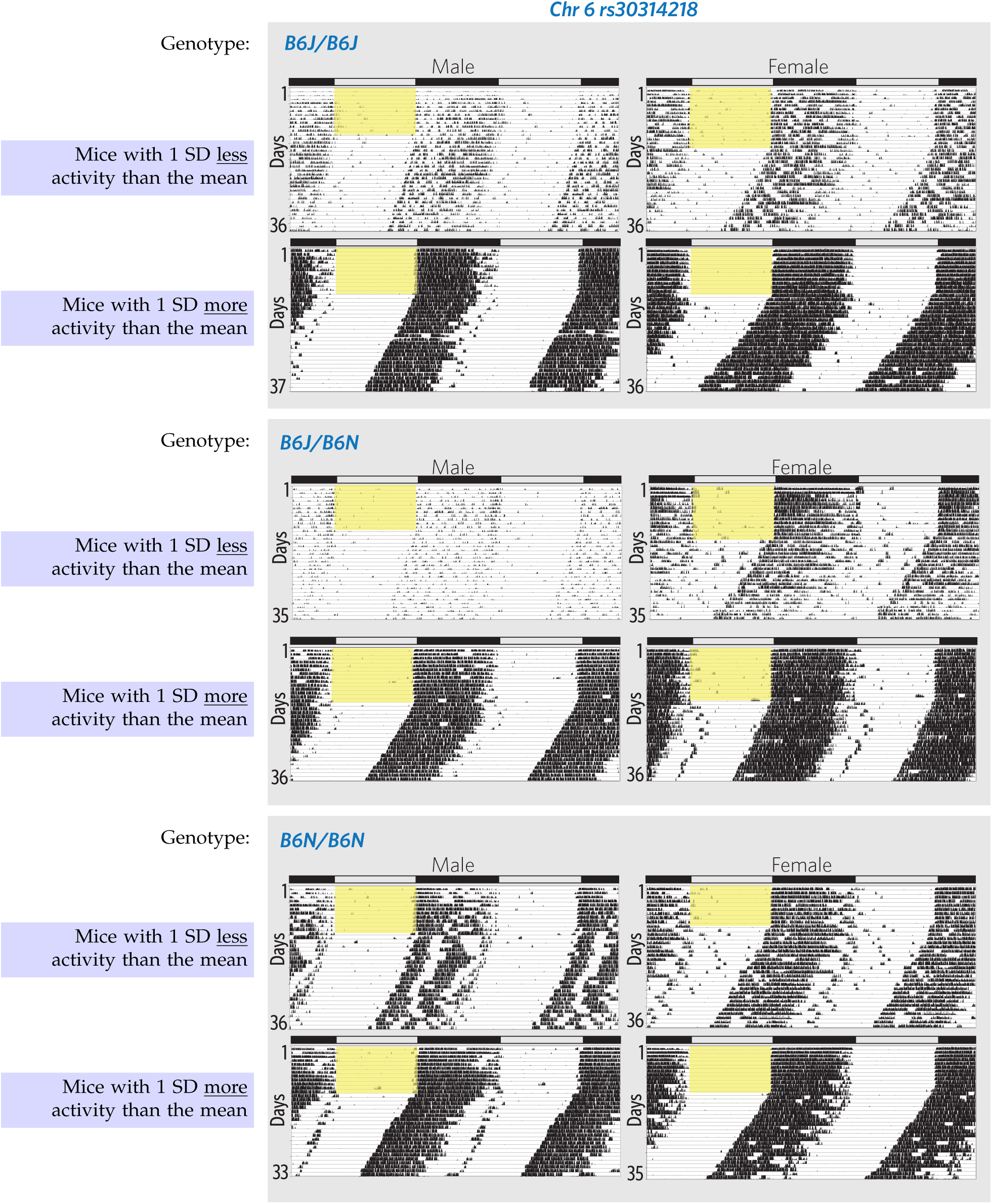
Double-plotted actograms illustrating phenotyping range of wheel running activity based on sex and genotype at rs30314218. The animals shown in this figure are marked in Figure 2 with larger circles. Yellow box indicates when lights were on. With groups defined by sex and genotype at rs30314218, each animal was chosen because it was approximately one group-specific SD above or below the group-specific mean. Note that there is much less variation in activity amongst the C57BL/6N homozygotes than amongst the other groups. The ID’s of the plotted mice are listed in the supplement.

**Figure 4.**
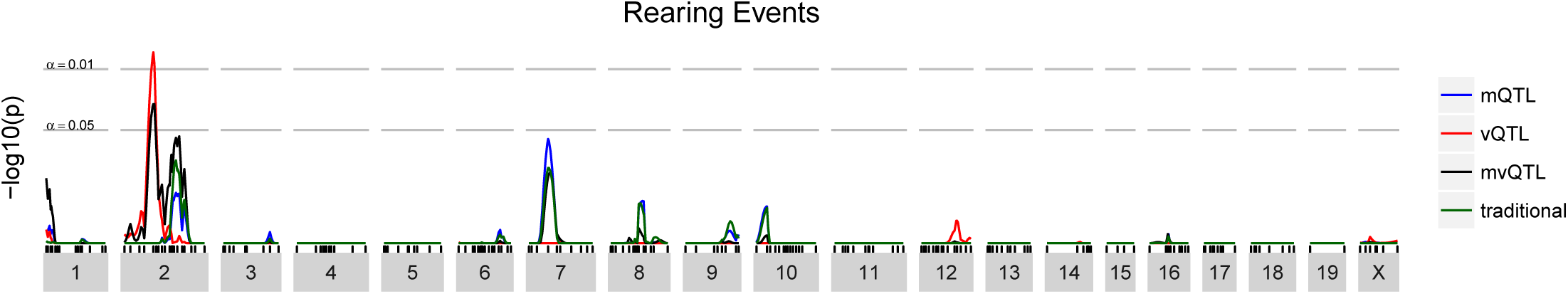
Genome scan for Bailey et al. rearing behavior. The x axis shows chromosomal location and the y axis shows FWER-controlling *p*-values for the association between each genomic locus and the Box-Cox transformed rearing behavior.

There are well-known and well-founded concerns that inappropriate scaling of phenotypes can produce spurious vQTL (Rönnegård and Valdar 2012; Sun *et al.* 2013; Shen and Ronnegard 2013). Therefore, the rearing phenotype was analyzed under a variety of additional transforms: none, log, square root, and 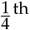 power (the transformation recommended by the Box-Cox procedure). Because the trait is a “count” and a positive mean-variance correlation was observed, the trait was further analyzed with a Poisson double generalized linear model with its canonical link function (log). In all cases, the same genomic region on chromosome 2 was identified as a statistically significant vQTL (*p* < 0.01) (Figure S6, Figure S8, Figure S7, and Figure S10). Though all transformations yielded similar results, we highlight the Box-Cox transformed analysis recommended for transformation selection in Rönnegård and Valdar (2011).

### Understanding the Novel QTL

In this case, the DGLM-based analysis identified a vQTL, a pattern of variation across genotypes not targeted by traditional, SLM-based, QTL analysis. The phenotype values, when stratified by genotype at the top locus, illustrate clear variance heterogeneity (Figure 5a). The effects and their standard errors estimated by the DGLM fitted at the top locus corroborate the impression from simply viewing the data, that the locus is a vQTL but not an mQTL (Figure 5b).

**Figure 5.**
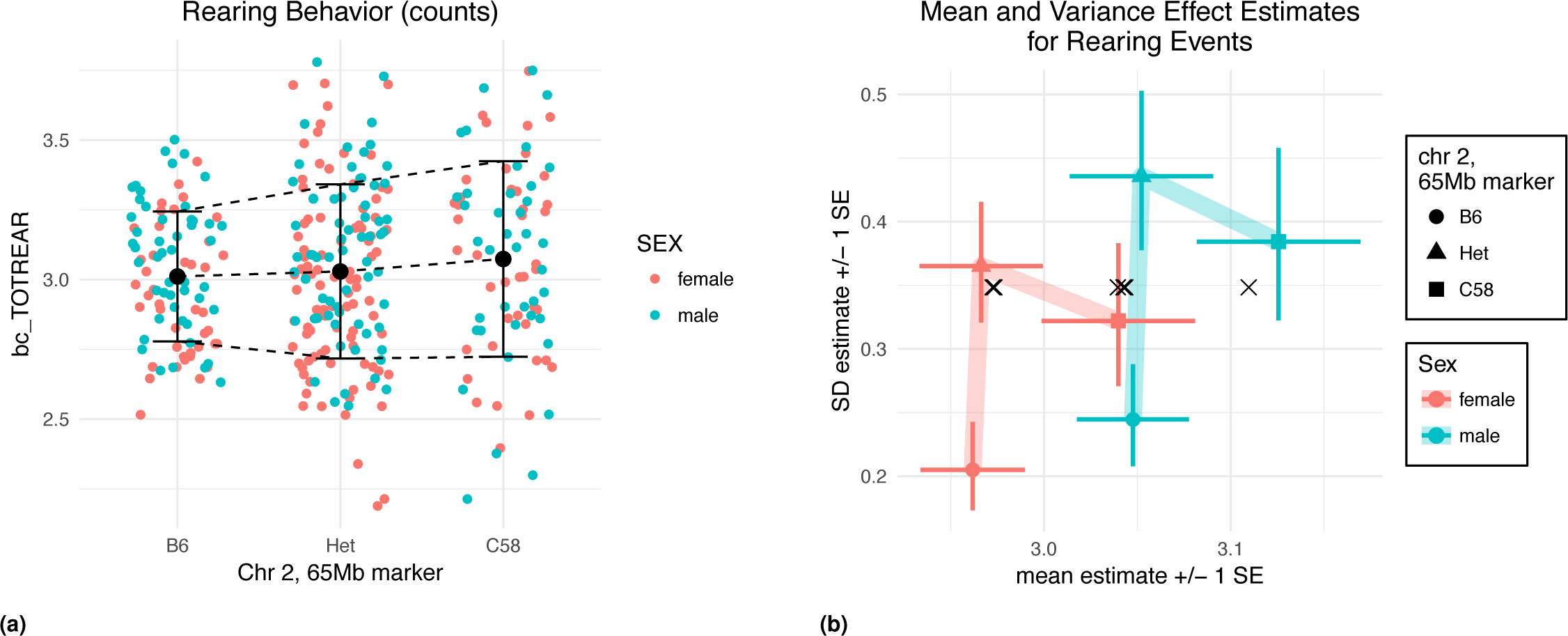
(a) “Total Rearing Events”, transformed by the Box-Cox procedure, stratified by sex and genotype at the top marker. (b) Predicted mean and variance of mice according to sex and allele at the top marker.

## DISCUSSION

We have demonstrated through two case studies that mean-variance QTL mapping based on the DGLM meaningfully expands the scope of QTL that can be detected to include both vQTL and more mQTL at loci that exhibit variance heterogeneity. In an era where ever more complete and complex data on biological systems is becoming available, this modest elaboration of an existing approach represents a single step toward the broader goal of characterizing the wide array of patterns of association between genotype, environment, and phenotype.

In the reanalysis of Kumar et al., mean-variance QTL mapping identified the same QTL as traditional, SLM-based QTL mapping for cocaine response traits and one novel mQTL for a circadian behavior trait. Such an mQTL would likely have been detected by a traditional QTL analysis with a larger mapping population: Through simulation, we estimated that the additional power to detect the mQTL was equivalent to the power increase that would have come from increasing the sample size by *≈*100 mice, to *≈*350. Given the considerable effort and expense associated with conducting an experimental cross or expanding the size of the mapping population, there seems to be little to be gained by omitting a DGLM-based analysis.

In the reanalysis of Bailey et al., mean-variance QTL mapping identified a novel vQTL for an exploratory behavior. A vQTL such as this would not be detected by the traditional QTL analysis no matter how large the mapping population because the pattern is entirely undetectable by the SLM.

The identification of a vQTL raises important issues related to phenotype transformation and the interpretation of findings, but both are manageable, as we have illustrated here. The criticism that a spurious vQTL can arise as the result of an inappropriate transformation is based on the observation that when genotype means are unequal, there always exists a (potentially exotic) transformation that diminishes the extent of variance heterogeneity (Sun *et al.* 2013). Thus, any other transformation (including none at all) can be seen as inflationary toward variance heterogeneity. In this context, however, an “inappropriate transformation” leads not to the misclassification of a non-QTL as a QTL, but an mQTL as a vQTL.

To the extent that the goal of QTL mapping is to understand the genetic architecture of a trait, this criticism is valid and should be addressed by considering a wide range of transformations, alternative models, and parameterizations. To the extent that the goal of QTL mapping is to identify genomic regions that contain genes and regulatory factors that influence a trait, we argue that such a misclassification is largely irrelevant. Whether we pursue bioinformatic follow-up to identify QTN in a region because it was identified as an mQTL or a vQTL need not change our downstream efforts.

In summary, we advocate for the use of mean-variance QTL mapping not as an additional flourish to consider after conducting an SLM-based QTL mapping effort, but rather as a drop-in replacement. This approach should not be too alien — when variance heterogeneity is absent, it simplifies to the well-known SLM-based approach. Full-featured software that implements this approach is described in a companion article (Corty and Valdar 2018+ [r/vqtl]).

Lastly, we note an additional benefit conferred by mean-variance QTL mapping not discussed in depth here. Variance heterogeneity can also derive from factors acting in the “background”, that is, arising from experimental or biological variables that are outside the main focus of testing but that nonetheless predict phenotypic variability and thereby inform the relative precision of one individual’s phenotype over another. In the case studies presented here, the only background factor considered was sex. But, more generally, any factor that a researcher considers as a potentially important covariate that should be modeled can be included not only as a mean covariate (as with the SLM) but also as a variance covariate. In a companion article, we describe how accommodating such background factors can deliver additional power to detect mQTL, vQTL, and mvQTL (Corty and Valdar 2018+ [BVH]).

## ACKNOWLEDGMENTS

This work was primarily funded by a National Institutes of General Medical Sciences (NIGMS) grant to WV (R01-GM104125) and a National Institutes of Mental Health grant to RWC (F30-MH108265). RWC received additional support from NIGMS grant T32-GM067553 and National Library of Medicine grant T32-LM012420.

## SUPPLEMENTARY MATERIALS

### Supplement to Kumar Reanalysis

**Figure S1.**
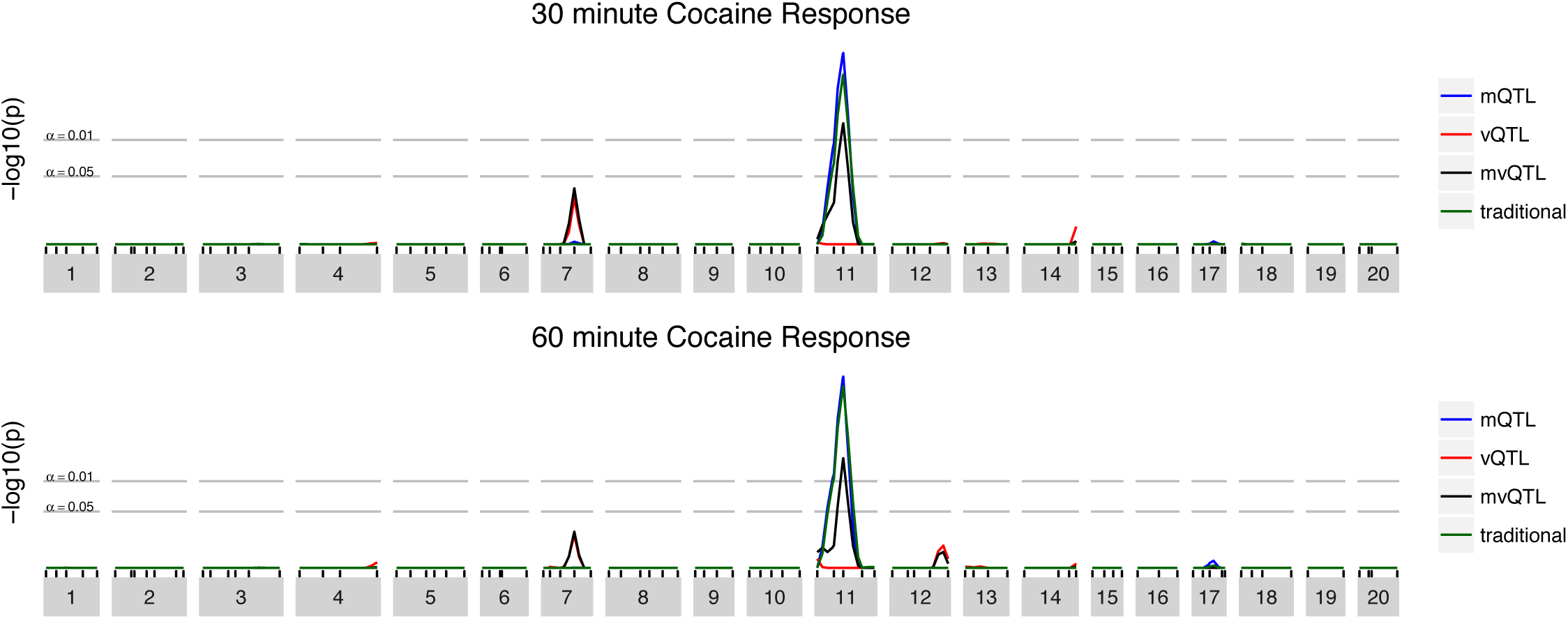
Replicated scans from Kumar *et al.* (2013)

**Figure S2.**
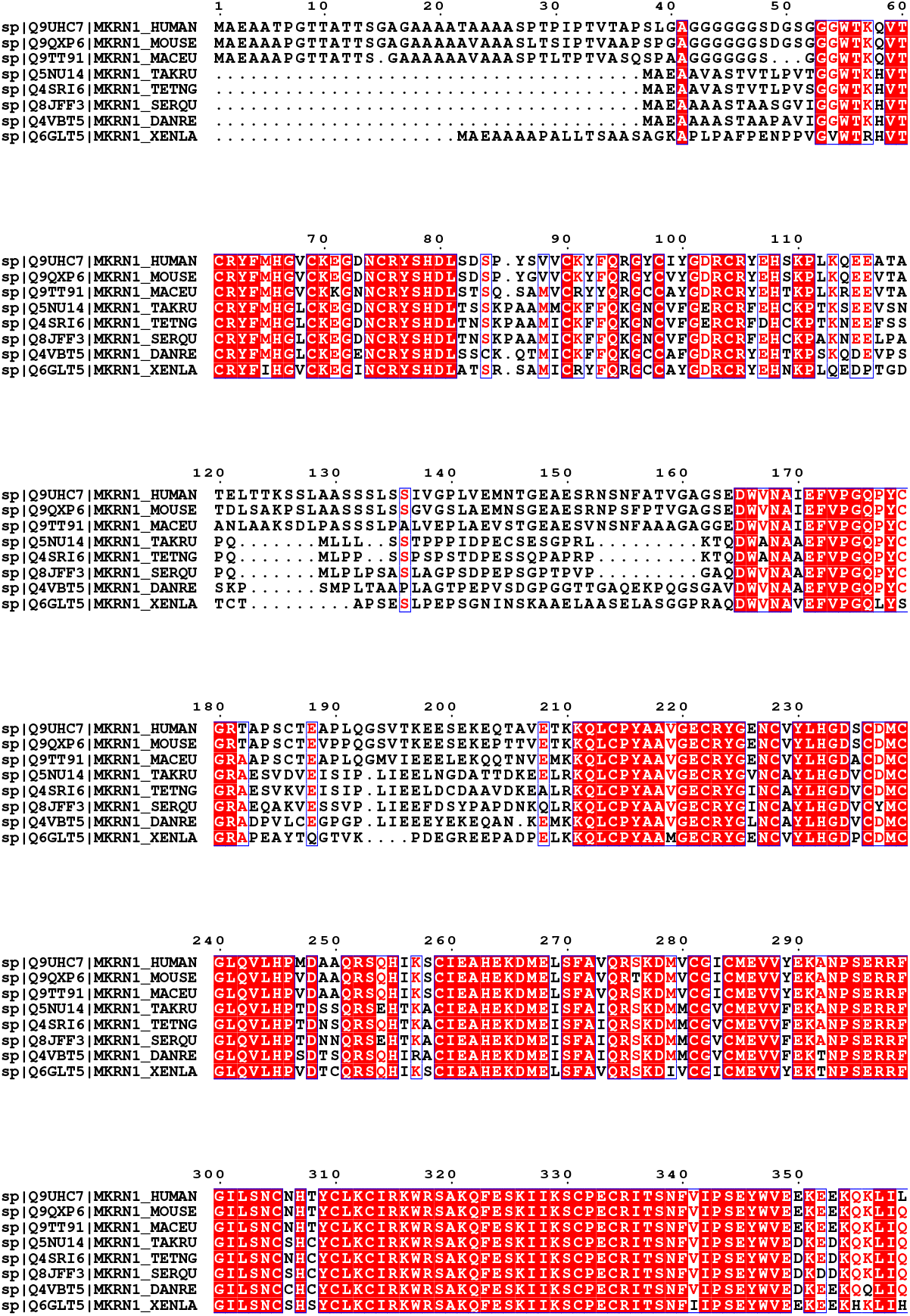
Page one of *Mkrn1* alignment. Note that the amino acid at position 346 is conserved across all species. See next page for species labels.

**Figure S3.**
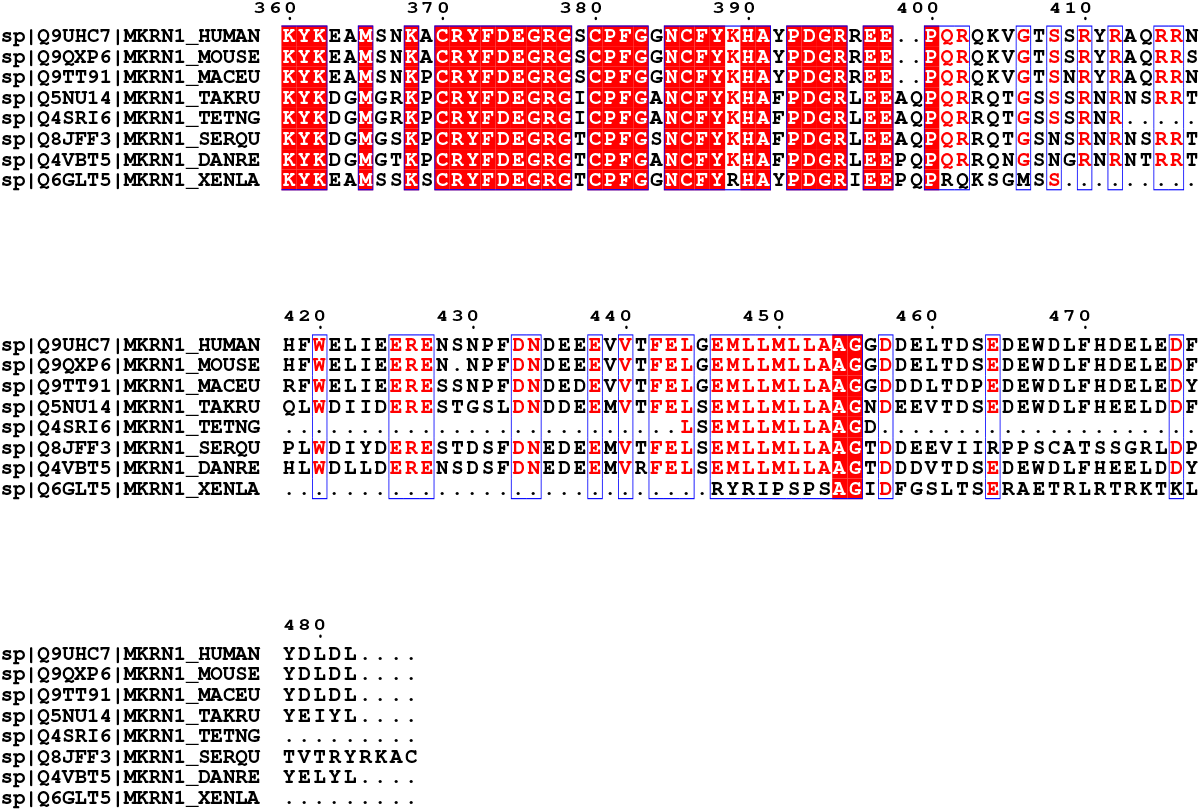
Page two of *Mkrn1* alignment.

**Figure S4.**
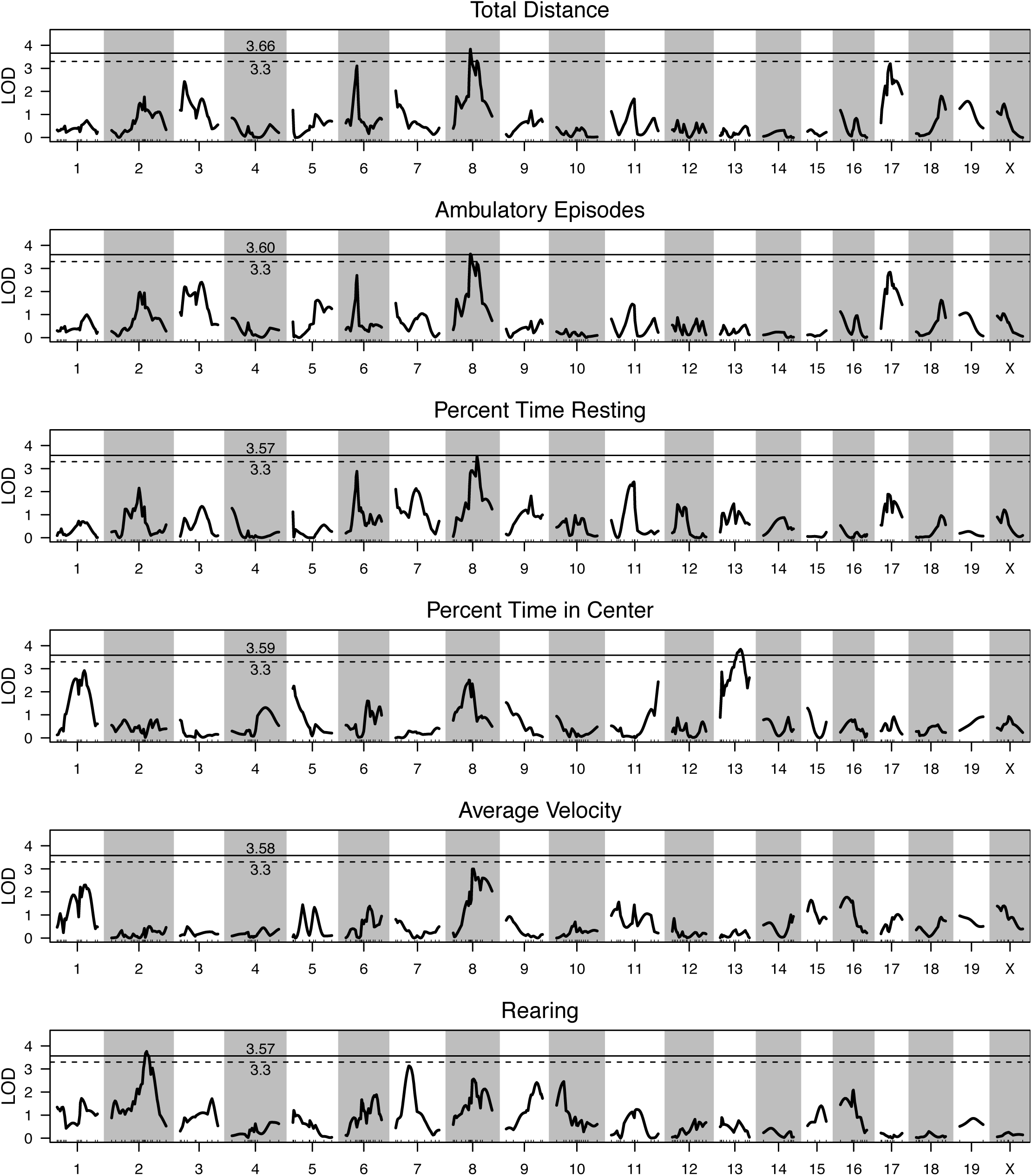
Replication of genome scans from original Bailey analysis. LOD curves are visually identical to originally-published LOD curves, but thresholds, estimated based on the described methods, are meaningfully higher.

**Figure S5.**
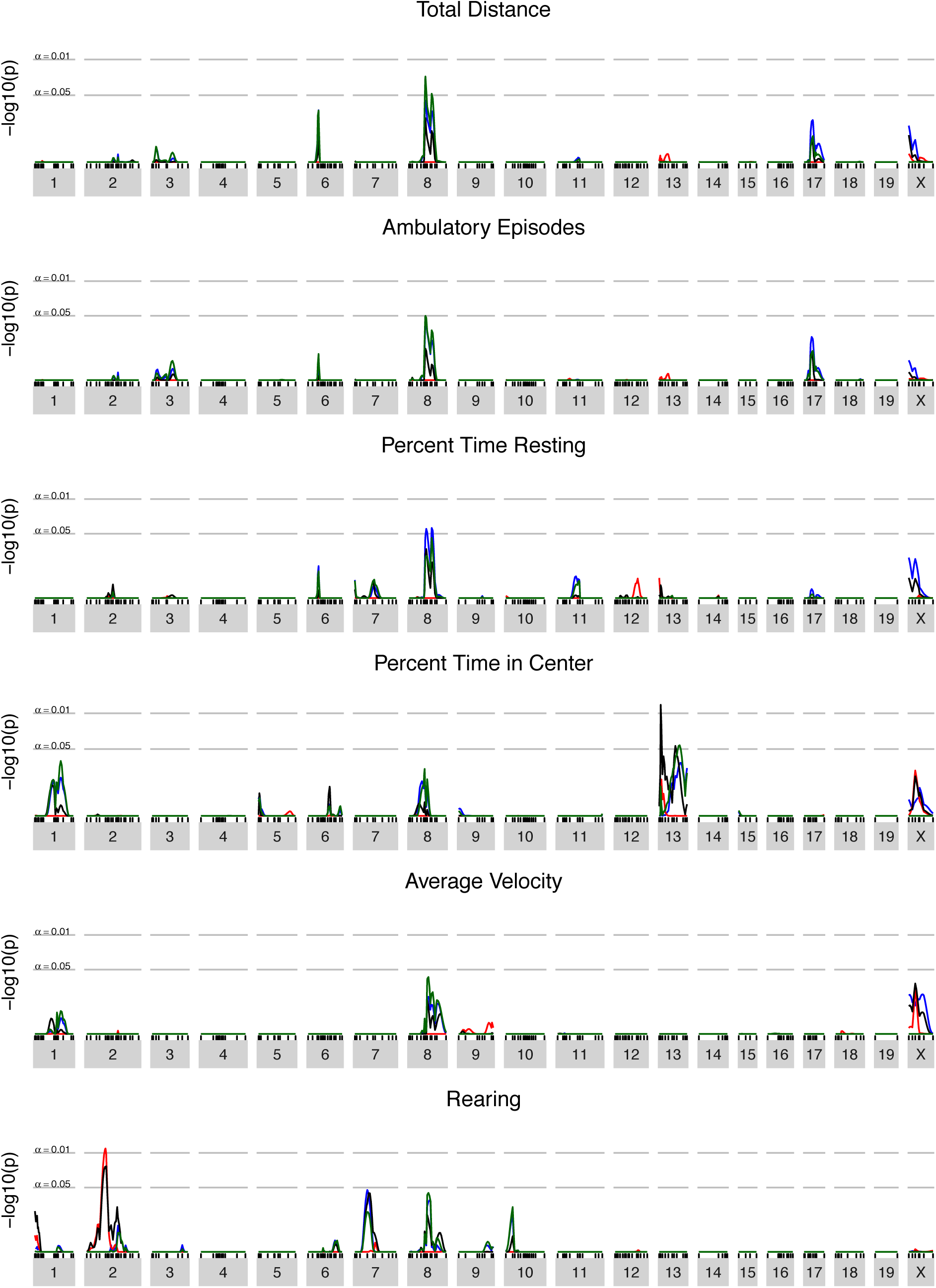
DGLM-based reanalysis of all traits measured in Bailey et al., all transformed by the rank-based inverse normal transform.

**Figure S6.**
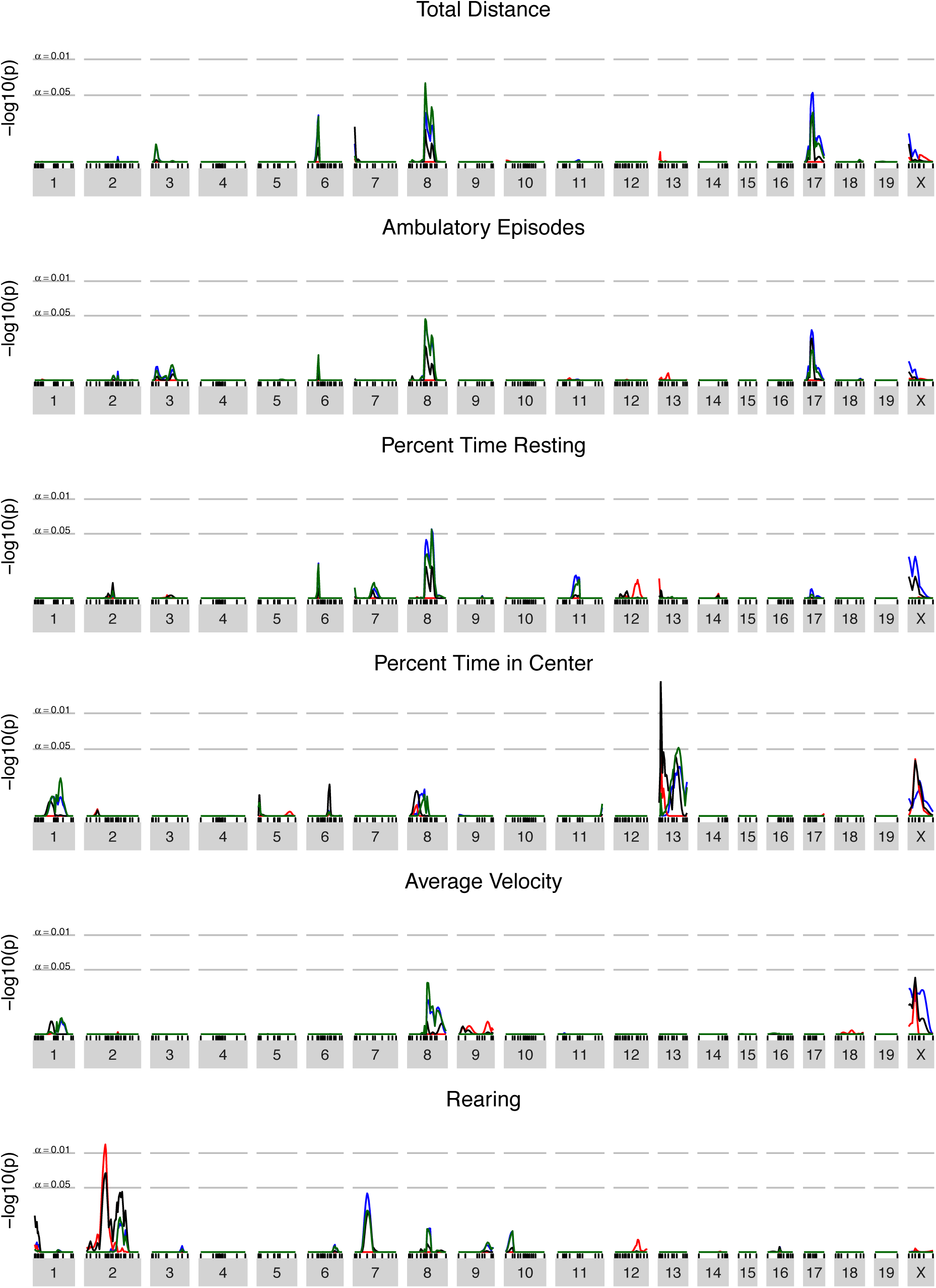
DGLM-based reanalysis of all traits measured in Bailey et al., all transformed by the Box-Cox transform. Box-Cox exponents were 1, 1, 0, 0.75, 0, 0.25, respectively.

**Figure S7.**
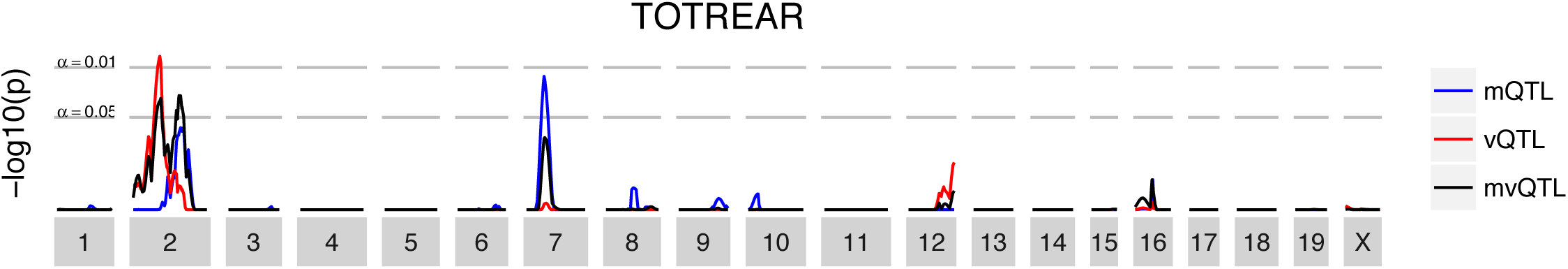
“Rearing” trait analyzed with no transformation.

**Figure S8.**
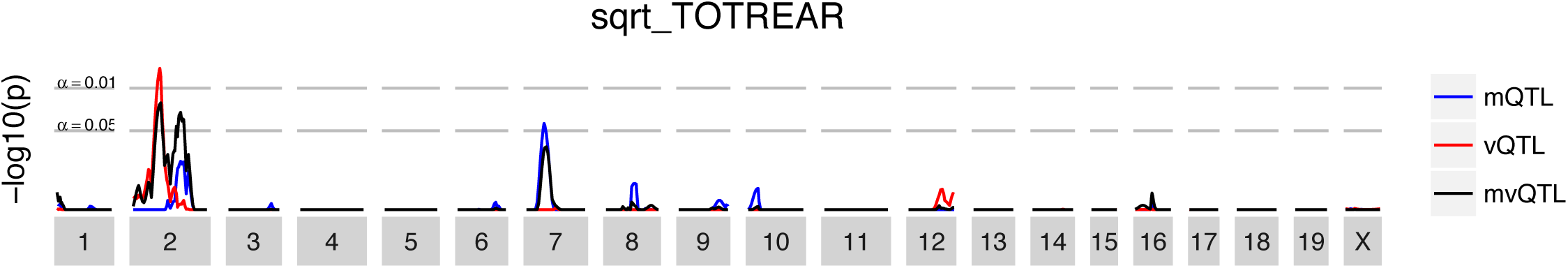
“Rearing” trait analyzed with square root transformation.

**Figure S9.**
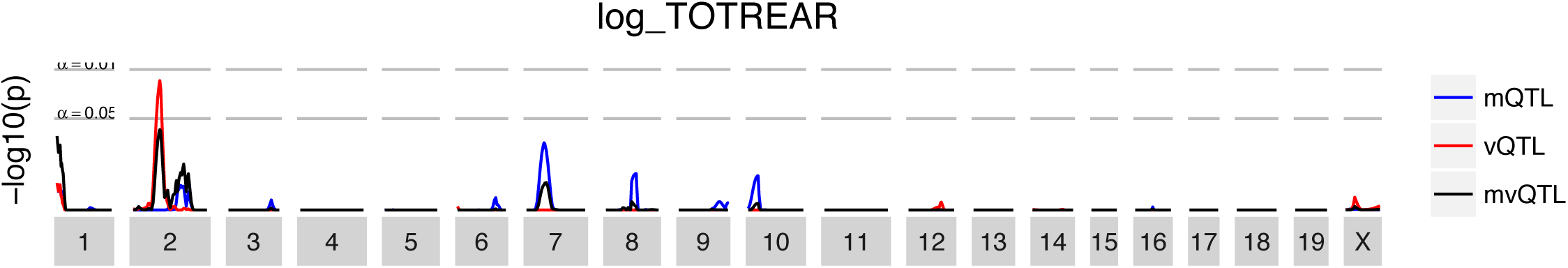
“Rearing” trait analyzed with log transformation.

**Figure S10.**
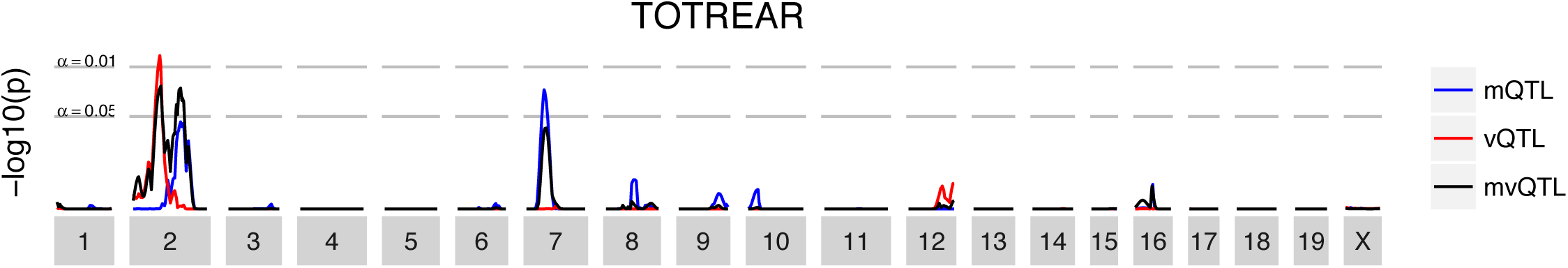
“Rearing” trait analyzed with Poisson generalized linear model.

## LITERATURE CITED

Allen Institute for Brain Science, 2015 Allen Mouse Brain Atlas. Allen Mouse Brain Atlas 2.

Bailey, J. S., L. Grabowski-Boase, B. M. Steffy, T. Wiltshire, G. A. Churchill, et al., 2008 Identification of quantitative trait loci for locomotor activation and anxiety using closely related inbred strains. Genes. Brain. Behav. 7: 761–9.

Banks, G., I. Heise, B. Starbuck, T. Osborne, L. Wisby, et al., 2015 Genetic background influences age-related decline in visual and nonvisual retinal responses, circadian rhythms, and sleep. Neurobiol. Aging 36: 380–393.

Bogue, M. a., G. a. Churchill, and E. J. Chesler, 2015 Collaborative Cross and Diversity Outbred data resources in the Mouse Phenome Database. Mamm. Genome 1.

Broman, K. W. and Ś. Sen, 2009 Single QTL Analysis. In A Guid. To QTL Mapp. with R/qtl, pp. 75–133.

Cao, Y., P. Wei, M. Bailey, J. S. K. Kauwe, and T. J. Maxwell, 2014 A versatile omnibus test for detecting mean and variance heterogeneity. Genet. Epidemiol. 38: 51–59.

Churchill, G. A. and R. W. Doerge, 1994 Empirical Threshold Values for Quantitative Trait Mapping. Genetics 138: 963–971.

Cox, A., C. L. Ackert-Bicknell, B. L. Dumont, Y. Ding, J. T. Bell, et al., 2009 A New Standard Genetic Map for the Laboratory Mouse. Genetics 182: 1335–1344.

Dudbridge, F. and B. P. C. Koeleman, 2004 Efficient computation of significance levels for multiple associations in large studies of correlated data, including genomewide association studies. Am. J. Hum. Genet. 75: 424–35.

Forsberg, S. K. G., M. E. Andreatta, X.-Y. Huang, J. Danku, D. E. Salt, et al., 2015 The Multi-allelic Genetic Architecture of a Variance-heterogeneity Locus for Molybdenum Accumulation Acts as a Source of Unexplained Additive Genetic Variance. bioRxiv p. 019323.

Geiler-Samerotte, K. A., C. R. Bauer, S. Li, N. Ziv, D. Gresham, et al., 2013 The details in the distributions: Why and how to study phenotypic variability. Curr. Opin. Biotechnol. 24: 752–759.

Haley, C. S. and S. a. Knott, 1992 A simple regression method for mapping quantitative trait loci in line crosses using flanking markers. Heredity (Edinb). 69: 315–24.

Kapushesky, M., I. Emam, E. Holloway, P. Kurnosov, A. Zorin, et al., 2009 Gene expression Atlas at the European Bioinformatics Institute. Nucleic Acids Res. 38.

Keane, T. M., L. Goodstadt, P. Danecek, M. A. White, K. Wong, et al., 2011 Mouse genomic variation and its effect on phenotypes and gene regulation. Nature 477: 289–294.

Kim, J. H., S. M. Park, M. R. Kang, S. Y. Oh, T. H. Lee, et al., 2005 Ubiquitin ligase MKRN1 modulates telomere length homeostasis through a proteolysis of hTERT. Genes Dev. 19: 776–781.

Kumar, V., K. Kim, C. Joseph, S. Kourrich, S.-H. Yoo, et al., 2013 C57BL/6N Mutation in Cytoplasmic FMRP interacting protein 2 Regulates Cocaine Response. Science (80-.). 342: 1508–1512.

Lander, E. S. and S. Botstein, 1989 Mapping mendelian factors underlying quantitative traits using RFLP linkage maps. Genetics 121: 185.

Martínez, O. and R. N. Curnow, 1992 Estimating the locations and the sizes of the effects of quantitative trait loci using flanking markers. Theor. Appl. Genet. 85: 480–488.

McWilliam, H., W. Li, M. Uludag, S. Squizzato, Y. M. Park, et al., 2013 Analysis Tool Web Services from the EMBL-EBI. Nucleic Acids Res. 41.

Nelson, R. M., M. E. Pettersson, and Ö. Carlborg, 2013 A century after Fisher: Time for a new paradigm in quantitative genetics. Trends Genet. 29: 669–676.

Paré, G., N. R. Cook, P. M. Ridker, and D. I. Chasman, 2010 On the use of variance per genotype as a tool to identify quantitative trait interaction effects: A report from the women’s genome health study. PLoS Genet. 6: 1–10.

Rönnegård, L. and W. Valdar, 2011 Detecting major genetic loci controlling phenotypic variability in experimental crosses. Genetics 188: 435–447.

Rönnegård, L. and W. Valdar, 2012 Recent developments in statistical methods for detecting genetic loci affecting phenotypic variability. BMC Genet. 13: 63.

Shen, X., M. Pettersson, L. Ronnegard, and O. Carlborg, 2012 Inheritance beyond plain heritability: variance-controlling genes in Arabidopsis thaliana. PLoS Genet. 8: e1002839.

Shen, X. and L. Ronnegard, 2013 Issues with data transformation in genome-wide association studies for phenotypic variability. F1000Research 2: 200.

Shimomura, K., S. S. Low-Zeddies, D. P. King, T. D. Steeves, A. Whiteley, et al., 2001 Genome-wide epistatic interaction analysis reveals complex genetic determinants of circadian behavior in mice. Genome Res. 11: 959–80.

Simon, M. M., S. Greenaway, J. K. White, H. Fuchs, V. Gailus-Durner, et al., 2013 A comparative phenotypic and genomic analysis of C57BL/6J and C57BL/6N mouse strains. Genome Biol. 14.

Smyth, G. K., 1989 Generalized linear models with varying dispersion. J. R. Stat. Soc. Ser. B Methodol. 51: 47–60.

Sun, X., R. Elston, N. Morris, and X. Zhu, 2013 What is the significance of difference in phenotypic variability across SNP genotypes? Am. J. Hum. Genet. 93: 390–397.

Valdar, W., J. Flint, and R. Mott, 2006 Simulating the Collaborative Cross: power of quantitative trait loci detection and mapping resolution in large sets of recombinant inbred strains of mice. Genetics 172: 1783–1797.

